# Psilocybin Prolongs the Neurovascular Coupling Response in Mouse Visual Cortex

**DOI:** 10.1101/2025.07.25.666803

**Authors:** Rick T. Zirkel, Matthew Isaacson, Clara Liao, Matthew Yi, Keri Yamaguchi, Daniel Rivera, Amy Kuceyeski, Nozomi Nishimura, Alex C. Kwan, Chris B. Schaffer

**Affiliations:** Meinig School of Biomedical Engineering, Cornell University, Ithaca, NY, USA; Interdepartmental Neuroscience Program, Yale University, New Haven, CT, USA; Department of Radiology, Weill Cornell Medicine, New York, NY, USA; Department of Psychiatry, Weill Cornell Medicine, New York, NY, USA

## Abstract

Psilocybin has profound therapeutic potential for various mental health disorders, but its mechanisms of action are unknown. Functional MRI studies have reported the effects of psilocybin on brain activity and connectivity; however, these measurements rely on neurovascular coupling to infer neural activity changes and assume that blood flow responses to neural activity are not altered by psilocybin. Using two-photon excited fluorescence imaging in the visual cortex of awake mice to simultaneously measure neural activity and capillary blood flow dynamics, we found that psilocybin administration prolonged the increase in visual stimulus-evoked capillary blood flow – an effect which was reduced by pretreatment with a 5-HT_2A_R antagonist – despite not causing changes in the stimulus-evoked neural response. Multi-modal widefield imaging also showed that psilocybin extends the stimulus-evoked vascular responses in surface vessels with no observed effect on the population neural response. Computational simulation with a whole-brain neural mass model showed that prolonged neurovascular coupling responses can lead to spurious increases in BOLD-based measures of functional connectivity. Together, these findings demonstrate that psilocybin broadens neurovascular responses in the brain and highlights the importance of accounting for these effects when interpreting human neuroimaging data of psychedelic drug action.

---

Psychedelics have shown remarkable therapeutic potential for the treatment of mental health disorders such as depression, addiction, and anxiety ^1–4^. This resurgence has spurred human studies to investigate the mechanisms of psychedelic drug action, mainly with the use of functional magnetic resonance imaging (fMRI) to assess brain activity ^5–9^. Many fMRI studies investigating psychedelics have reported alterations in brain activity and connectivity patterns including increased ^5,6,8^ or decreased ^5,8,9^ global functional connectivity, reduced default mode network integrity ^5,6,8,9^, and heightened entropy ^9^ in brain signals. Such findings have shaped current models of how psychedelics affect cognition and consciousness.

fMRI relies on the principle of neurovascular coupling (NVC), in which neural activity changes are accompanied by regional changes in blood flow ^10^, which enables the measurement of blood-oxygenation-level-dependent (BOLD) signals as a proxy for neural activity ^11–14^. Although increases in blood flow are associated with increased neural activity, the relationship between the BOLD signal and neural activity is a complex process. BOLD depends on the rates of oxygen metabolism as well as the regulation of blood flow through vessel diameter changes that involve multiple signaling pathways and cell types ^15–19^, and these processes are not always perfectly maintained. Various conditions, such as oxidative stress, inflammation, endothelial dysfunction, and altered neuron-astrocyte signaling can disrupt this relationship ^20–23^. In Alzheimer’s disease, elevated reactive oxygen species damage vascular cells, impairing signaling between neurons, astrocytes, and blood vessels ^21,23^. In stroke and ischemic injury, astrocyte dysfunction and gliosis compromise the ability to regulate blood flow ^20,22^. In these cases, BOLD signals may no longer reflect neural activity in the same way as in healthy states – often referred to as neurovascular uncoupling ^24^ – complicating fMRI interpretation.

Psychedelic drugs may also alter the relationship between neural activity and associated changes in blood flow. Psychedelics are similar in structure to serotonin (5-HT) and have a high binding affinity for various serotonin receptors ^25^, including 5-HT_2A_R which is the primary receptor implicated in the hallucinogenic effects of psychedelics ^26,27^. Several serotonin receptors (e.g. 5-HT _1A_, _1B_, _2A_, _2B_) are expressed in cell types implicated in NVC ^28^. Activation of these receptors can cause the release of vasodilatory substances from endothelial cells, as well as contribute to vasoconstriction in cerebral arteries. Recent studies have demonstrated that psilocybin, and its receptor agonism at the 5-HT_2A_R, induce changes in cerebral blood flow and acute constriction of the internal carotid artery diameter in healthy humans ^29–31^. Serotonin itself is a vasoactive molecule ^32^, and can have both vasoconstrictive and vasodilatory effects in a context and region specific manner ^33–35^. Thus, the serotonin receptor interactions of psychedelics may directly alter blood flow responses to neural activity, independent of any changes in that neural activity ^36–38^.

In this study, we used multiple imaging modalities to simultaneously measure neural activity and vascular dynamics (blood flow velocity, vessel dilation, and blood oxygenation) in response to a visual stimulus before and after treatment with psilocybin. Two-photon excited fluorescence microscopy (2PEF) and wide field multimodal imaging were used to characterize stimulus-induced neurovascular responses in the visual cortex at the microscale and mesoscale, respectively. Pretreatment with MDL100907, a highly selective 5-HT_2A_ receptor antagonist, allowed us to assess the involvement of 5-HT_2A_R in NVC modulation. Based on our findings, we then simulated how the psilocybin-induced alterations in NVC could affect downstream BOLD measurements in fMRI under identical neural activity profiles, potentially leading to misinterpretation of brain functional connectivity changes in human psilocybin research.

## Results

### Visual stimulus-evoked capillary blood flow, but not neural activity, was prolonged by psilocybin

Using in vivo 2PEF imaging in awake, head-fixed mice, we measured calcium transients in individual GCaMP6f-expressing neurons alongside capillary blood flow and diameter changes in the visual cortex in response to a visual stimulus. During each trial, mice were shown a repeated four-direction pattern drifting grating stimulus (Fig. 1a). Using fast (167 Hz) arbitrary line scanning, neural activity response, capillary flow velocity, and capillary diameter were simultaneously recorded in V1 layer II/III (Fig. 1b-d). Imaging was repeated in the same neurons and capillaries before and after treatment with either psilocybin (1 mg/kg, I.P.) or saline, allowing us to assess treatment-induced changes in neurovascular coupling within the same cells and vessels.

**Fig. 1.**
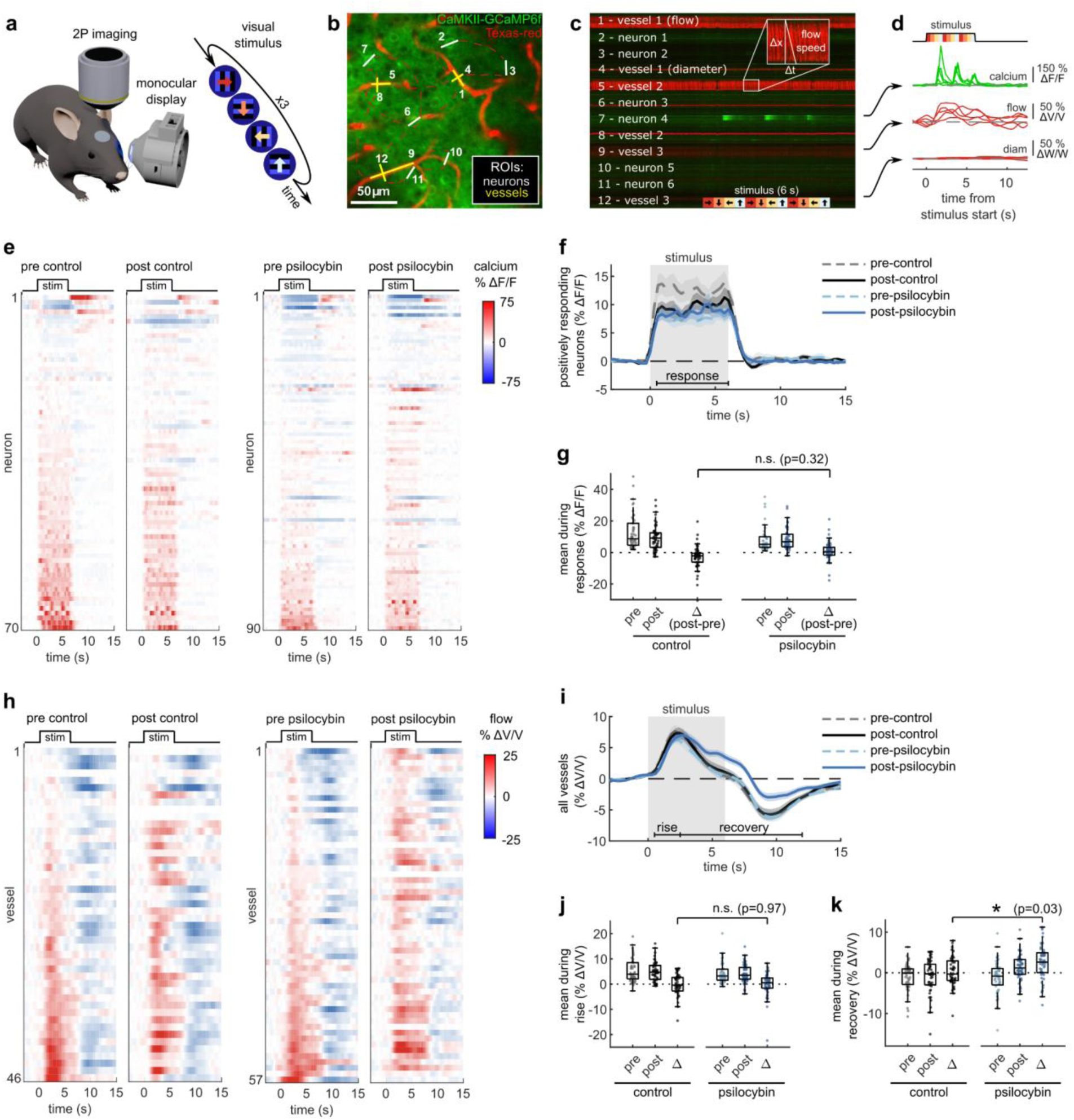
Effects of psilocybin on neural and capillary blood flow response in visual cortex to a visual stimulus. **a:** Two photon excited fluorescence imaging of V1 in an awake mouse during monocular visual stimulation with a 4-direction repeated drifting grating. **b:** Arbitrary line scan path recording simultaneous calcium activity from neurons (green channel, traced with white line ROIs) and diameter and blood flow velocity in capillaries (red channel, with yellow line ROIs). **c:** Raw 2-channel imaging data from line scan path drawn in (b), with all ROIs labelled. The inset illustrates the calculation of vessel flow velocity from angled shadows produced by flowing unlabeled red blood cells. **d:** Example traces of calcium indicator fluorescence, vessel flow velocity, and vessel diameter changes (ROIs 7, 9, and 12 in (c)) during five repetitions of the visual stimulus. **e:** Raster plots of calcium indicator fluorescence change during the visual stimulus before and after control (left) or psilocybin treatment (right). **f:** Mean fluorescence change of all neurons classified as positively responding for each treatment condition. **g:** Box plots of mean fluorescence change during the stimulated time period (0.5-6 s) for each positively responding neuron in all conditions, with the mean post-pre difference values (Δ) shown for each treatment. **h:** Raster plots similar to (e) of capillary flow velocity change during the visual stimulus before and after treatment. **i:** Mean velocity change of all vessels for each treatment condition. **j-k:** Box plots of mean velocity change during the (j) rise time period (0.5-2.5 s) and (k) recovery time period (2.5-12 s), as well as the post-pre difference values (Δ). All mean traces are shown with +/-S.E.M. in shaded regions. All box plots display median, 25th, and 75th percentiles, with whiskers extending to the smallest/largest non-outliers. Significant effects of the treatment were determined with linear-mixed effects modeling followed by benjamini-hochberg correction for multiple comparisons (1 star: adj. p<0.05, 2 stars: adj. p<0.01).

We observed no consistent changes in the stimulus-induced neural response after treatment with psilocybin (Fig. 1e). Most neurons increase neural activity in response to the visual stimulus (Extended Data Fig. 1a), and the mean visually evoked response from these positively respondig neurons was not significantly affected by the treatment (Fig. 1f, g). In contrast, the stimulus-induced capillary blood flow response was prolonged in many vessels after treatment with psilocybin (Fig. 1h). Compared to the average neural response, the capillary blood flow response was more complex, featuring an initial rise and peak (henceforth referred to as the “rise” phase) followed by a reduction and undershoot after stimulus offset before returning to baseline (referred to as the “recovery” phase). Psilocybin prolonged the blood flow response to a stimulus, specifically inducing no significant change in average velocity during the rise phase but a significantly greater mean flow velocity during the recovery phase (Fig. 1i-k).

Capillary diameter was also calculated for many of the same vessels in each condition, though we observed no clear dilatory response in these small vessels with the stimulus (Extended Data Fig. 1b-c). We also did not detect any significant change in dilatory responsiveness, baseline flow velocity, or baseline capillary diameter after treatment (Extended Data Fig. 1d-i). Taken together, these findings indicate that psilocybin alters neurovascular coupling (NVC) at the microscale by prolonging the capillary blood flow response to stimulus-induced neural activity.

### Widefield imaging supports prolonged stimulus-induced neurovascular responses after psilocybin

To assess whether the prolonged vascular response caused by psilocybin was evident at the population level, we conducted widefield multimodal imaging to measure stimulus-induced neural activity along with blood flow speed, vessel dilation, and blood oxygenation in larger surface vessels (Fig. 2a-c). Widefield fluorescence of GCaMP6f was used to characterize neural activity, while laser-speckle contrast imaging was used to quantify relative changes in blood flow speed and 530/560 nm reflectance was used to measure relative changes in blood oxygenation produced by the visual stimulus (Fig. 2b).

**Fig. 2.**
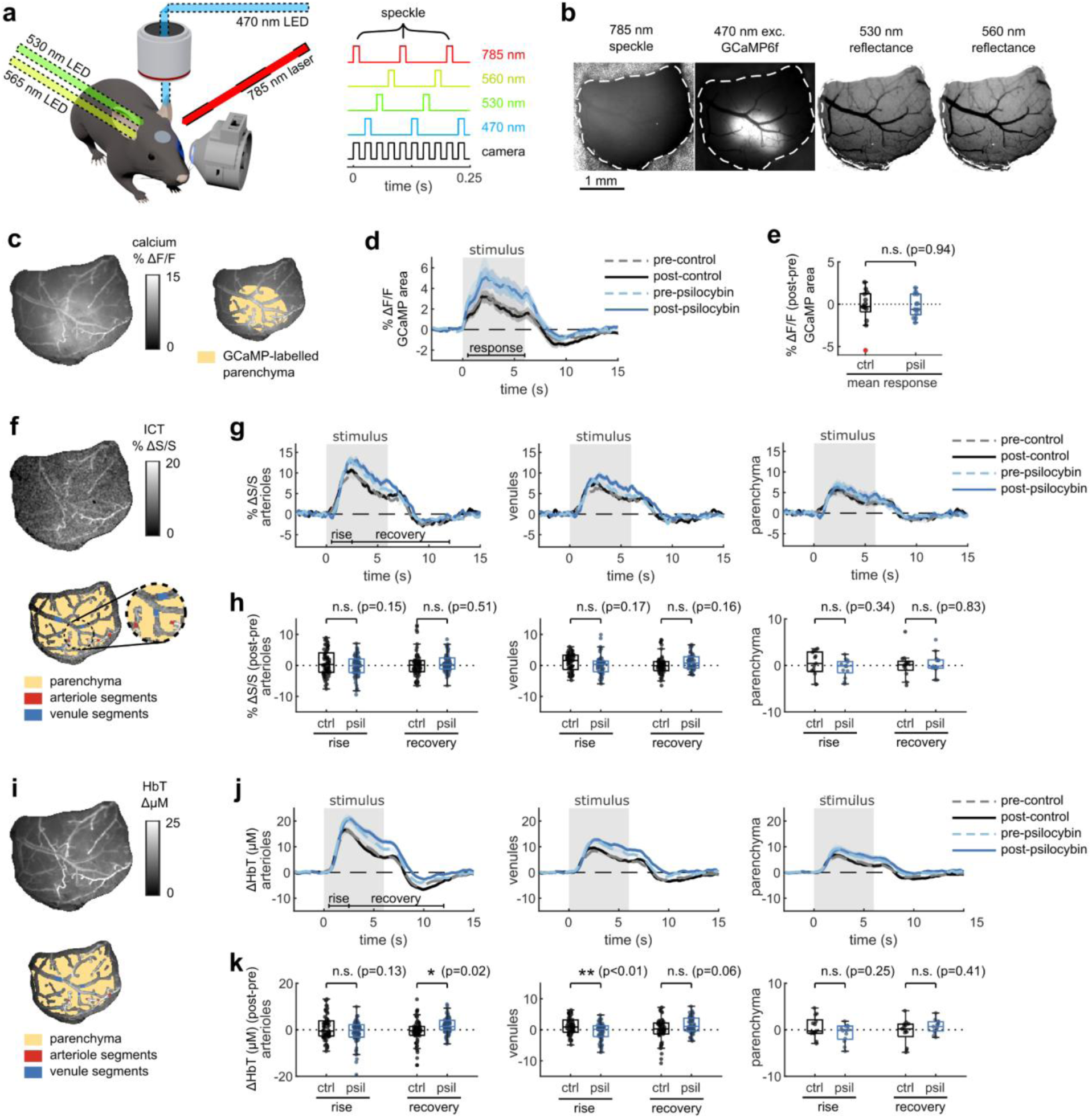
Multimodal widefield measurements of the effects of psilocybin on neural and vascular stimulus responses. **a:** (left) Overview of multimodal widefield imaging during monocular visual stimulation, with (right) four excitation light sources sequentially pulsed. **b:** Four-channel output of multimodal imaging: calcium indicator fluorescence (470-nm excitation), laser speckle contrast imaging for blood flow speed (785-nm LSCI), and multi-color reflectance for blood oxygenation determination (530- and 560-nm reflectance). **c:** (left) Example map of oxygen absorption-corrected calcium indicator fluorescence change after treatment, relative to before, averaged during the stimulus response time (0.5-6 s). (right) ROI drawn over the GCaMP6f-labelled parenchyma, excluding large vessels. **d:** Mean fluorescence change in GCaMP-labelled parenchyma for each treatment condition. **e:** Box plots of difference in mean fluorescence change during the stimulus response time period (0.5-6 s), with the mean post-pre difference (Δ) shown for each treatment. **f:** (top) Example map of normalized inverse correlation time (ICT) images calculated from speckle contrast imaging, averaged over the stimulus response time period (0.5-6 s). (bottom) ROIs drawn for whole-window parenchyma region, arteriole segments, and vessel segments. **g:** Mean relative speed changes over all manually-labelled arteriole segments (left), venule segments (middle), and parenchyma (right) for each treatment condition. **h:** Box plots of difference in mean speed change after treatment, relative to before, during the rise and recovery time periods for arteriole segments (left), venule segments (middle), and parenchyma (right). **i:** Example map of mean change in total hemoglobin (HbT) concentration during the stimulus response time period (0.5-6 s) (top), and illustration of ROIs (bottom). **j:** Mean relative HbT concentration changes in arteriole segments (left), venule segments (middle), and parenchyma (right). **k:** Box plots of mean HbT concentration change after treatment, relative to before, during the rise and recovery time periods for arteriole segments (left), venule segments (middle), and parenchyma (right). All mean traces are shown as mean +/-S.E.M. in shaded regions. All box plots display median, 25th, and 75th percentiles, with whiskers extending to the smallest/largest non-outliers. Significant effects of the treatment were determined with linear-mixed effects modeling followed by benjamini-hochberg correction for multiple comparisons (1 star: adj. p<0.05, 2 stars: adj. p<0.01).

With the same visual stimulus paradigm as before, we first calculated the average absorption-corrected relative fluorescence change in GCaMP6f-labelled V1 parenchyma, observing no noticeable change after treatment (Fig. 2c-e). However, relative blood flow speed responses in arteriole and venule segments trended toward the prolonged speed increases we observed in capillaries (Fig. 2f-g), though the increase in mean flow speed during the vessel recovery period was not statistically significant (Fig. 2h). The increase in total hemoglobin concentration after the stimulus also followed this trend, with a significant increase in arteriole total hemoglobin during the recovery phase after psilocybin (Fig. 2i-k). We also observed a significantly reduced increase in venule total hemoglobin during the rise phase after psilocybin. These changes in total hemoglobin response were driven by an increased deoxygenated hemoglobin response and a decreased, or potentially delayed, initial oxygenated hemoglobin response (Extended Data Fig. 2a-f). Finally, similar to that observed in microscale imaging, we observed no change in the vessel dilation response to the stimulus (Extended Data Fig. 2g-i).

The effects of psychedelics on neurovascular coupling may be drug-specific. In particular, a recent study described a shortened NVC response in mice after exposure to 2,5-dimethoxy-4-iodoamphetamine (DOI) ^37^, another hallucinogenic 5-HT_2A_R agonist. Therefore we repeated our widefield multimodal imaging protocol after treatment with DOI (10 mg/kg, I.P.) to determine if the discrepancy may be due to differing methods or to unique effects from these two drugs. We found that, similar to psilocybin, DOI did not change the neural response to the stimulus (Fig. 3a-b). However, DOI caused a significant reduction in the arteriole and venule dilation response (Fig. 3c, d). In agreement with the prior study, DOI appeared to cause a shortened blood flow response, resulting in a significant reduction in arteriole flow speed during the recovery phase (Fig. 3e, f). No significant shortening was observed in the total hemoglobin response (Fig. 3g, h), but a trend toward a shortened response was visible in the deoxygenated and oxygenated hemoglobin response, with a significantly larger deoxygenated hemoglobin response during the recovery phase in both arterioles and venules (Extended Data Fig. 3).

**Fig. 3.**
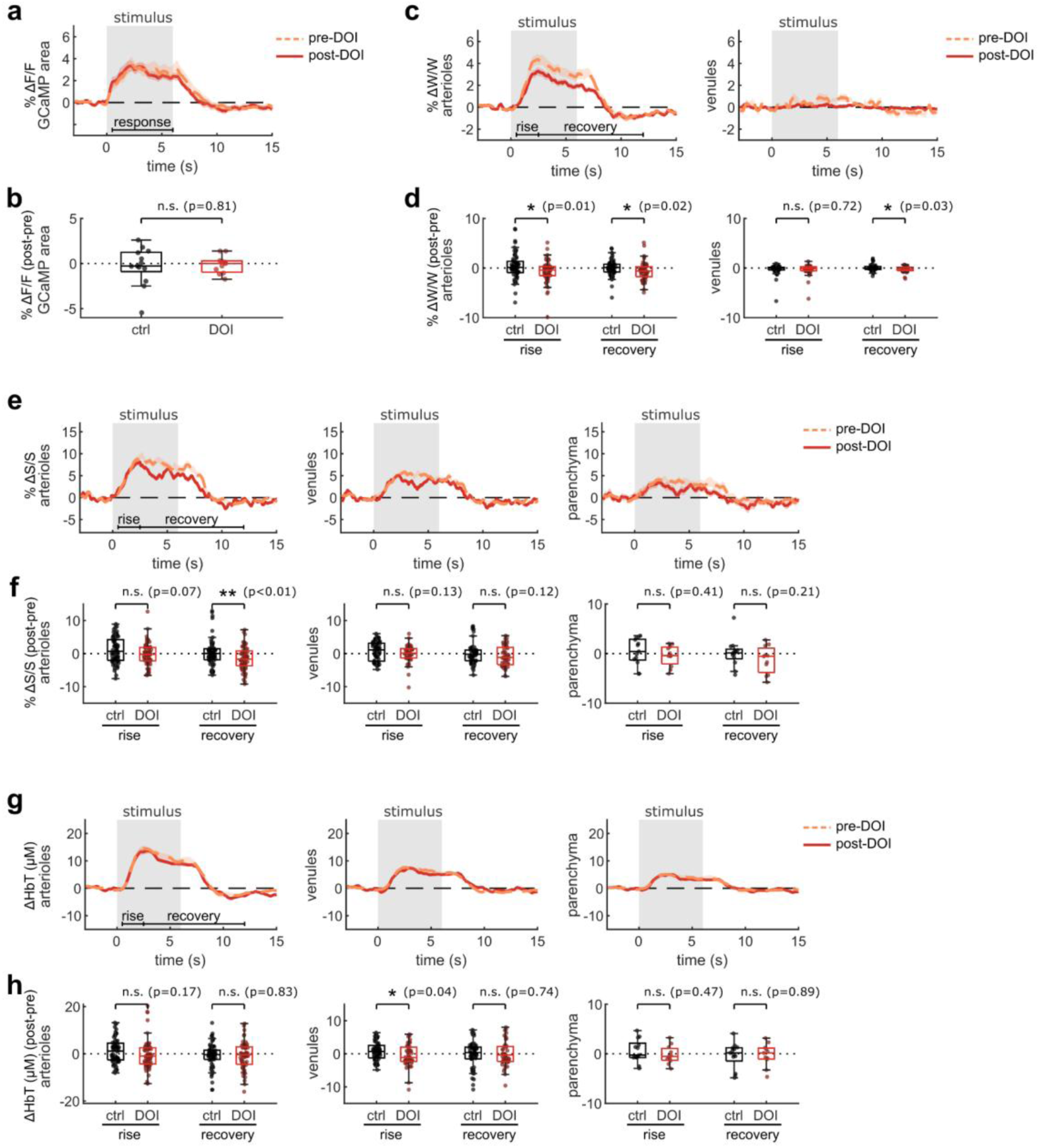
Multimodal widefield measurements after DOI treatment. **a:** Mean fluorescence change in GCaMP-labelled parenchyma region for each treatment condition. **b:** Box plots of difference in mean fluorescence change after treatment, relative to before, during the stimulus response time period (0.5-6 s) (control treatment data is replicated from Fig. 2d). **c:** Mean relative vessel width changes over all manually-labelled arteriole segments (left) and venule segments (right) for each treatment condition. **d:** Box plots of difference in mean speed change after treatment, relative to before, during the rise and recovery time periods for arteriole segments (left) and venule segments (right). **e:** Mean relative speed changes over all manually-labelled arteriole segments (left), venule segments (middle), and parenchyma (right) for each treatment condition. **f:** Box plots of difference in mean speed change after treatment, relative to before, during the rise and recovery time periods for arteriole segments (left), venule segments (middle), and parenchyma (right). **g:** Mean relative HbT concentration changes in arteriole segments (left), venule segments (middle), and parenchyma (right). **h:** Box plots of difference in mean HbT concentration after treatment, relative to before, during the response and recovery time periods for arteriole segments (left), venule segments (middle), and parenchyma (right). All mean traces are shown as mean +/-S.E.M. in shaded regions. All box plots display median, 25th, and 75th percentiles, with whiskers extending to the smallest/largest non-outliers. Data for the control treatment groups in all box plots are replicated from Figure 2. Significant effects of the treatment were determined with linear-mixed effects modeling followed by benjamini-hochberg correction for multiple comparisons (1 star: adj. p<0.05, 2 stars: adj. p<0.01).

### The effect of psilocybin on NVC was attenuated with a 5-HT_2A_R antagonist

To begin to understand the mechanism behind psilocybin-induced alteration of NVC, we re-examined microscale NVC changes during the administration of psilocybin after pretreatment with MDL100907 (referred to as MDL), a blocker of the 5-HT_2A_ receptor. Comparing our previously recorded measurements of average neural activity and vessel flow velocity response before and after psilocybin treatment, we found that pretreatment with MDL did not affect the stimulus-induced neural response (Fig. 4a-c), while the blood flow velocity response was relatively unchanged after psilocybin + MDL (Fig. 4d-g), in contrast to the significant effect caused by treatment with psilocybin alone (Fig 1). Thus, MDL partially attenuated the psilocybin-induced broadening of the NVC response. The mean flow response during the recovery phase was not significantly different from either the control or psilocybin treatments alone.

**Fig. 4.**
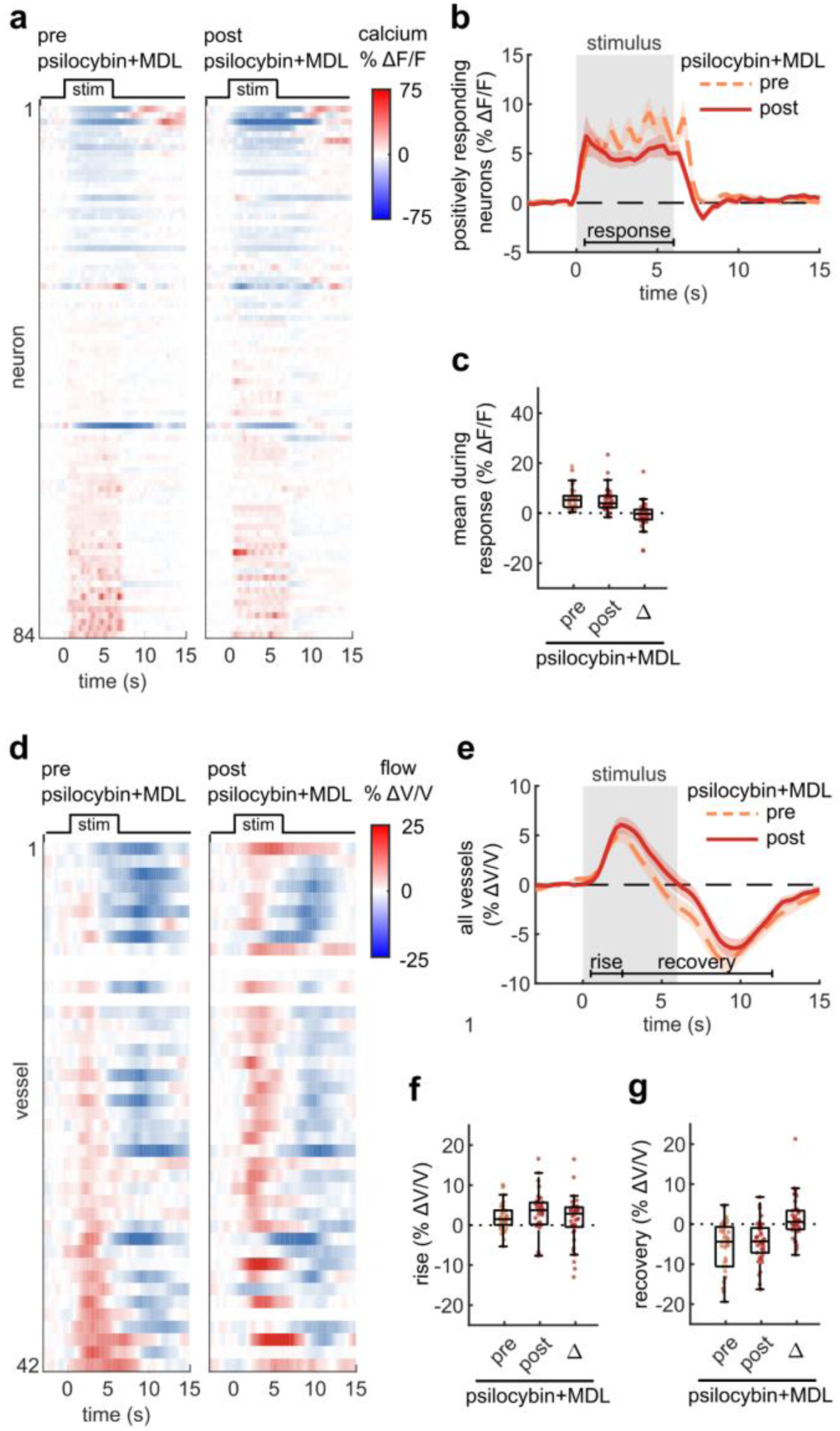
5HT_2A_R antagonist attenuated prolongation of NVC response due to psilocybin a: Raster plots of calcium indicator fluorescence change during the visual stimulus before and after psilocybin treatment in mice pretreated with MDL100907 (MDL). **b:** Mean fluorescence change of all neurons classified as positively responding for the pre and post treatment conditions. **c:** Box plots of mean fluorescence change during the stimulated time period (0.5-6 s) of each neuron in all conditions, with the mean post-pre difference (Δ) shown. **d:** Raster plots of vessel flow velocity change during the visual stimulus before and after psilocybin treatment in mice pretreated with MDL. **e:** Mean velocity change of all vessels for the pre (left) and post (right) psilocybin+MDL treatment. **f-g:** Box plots of mean velocity change during the (f) rise time period (0.5-2.5 s) and (g) recovery time period (2.5-12 s), as well as the post-pre Δ values. All mean traces are shown with +/-S.E.M. in shaded regions. All box plots display median, 25th, and 75th percentiles, with whiskers extending to the smallest/largest non-outliers.

### Psilocybin causes baseline blood vessel diameter increases

In both 2PEF and widefield imaging measurements, we did not observe any change in the stimulus-induced dilation response after psilocybin treatment. However, it is possible that psilocybin has effects on the baseline characteristics of the vasculature independent of the stimulus. We analyzed vessel diameters during the pre-stimulus time periods in our widefield imaging dataset, and observed a trend toward increased baseline vessel diameter in arterioles and a significant increase in venules after psilocybin treatment (Fig. 5a-b). This increase in baseline diameter was even greater after DOI treatment. While our measurement of baseline capillary diameter from line scans across a limited number of vessels did not show increased baseline diameter (Extended Data Fig. 2i), measurement of baseline capillary diameters from 2PEF volumetric imaging enables hundreds of more precise measurements. From this dataset, we observed a small but significant increase in capillary diameter after psilocybin (Fig. 4c-d). Additionally, psilocybin with MDL pretreatment did not result in a statistically significant increase in baseline diameter over the control treatment.

**Fig. 5.**
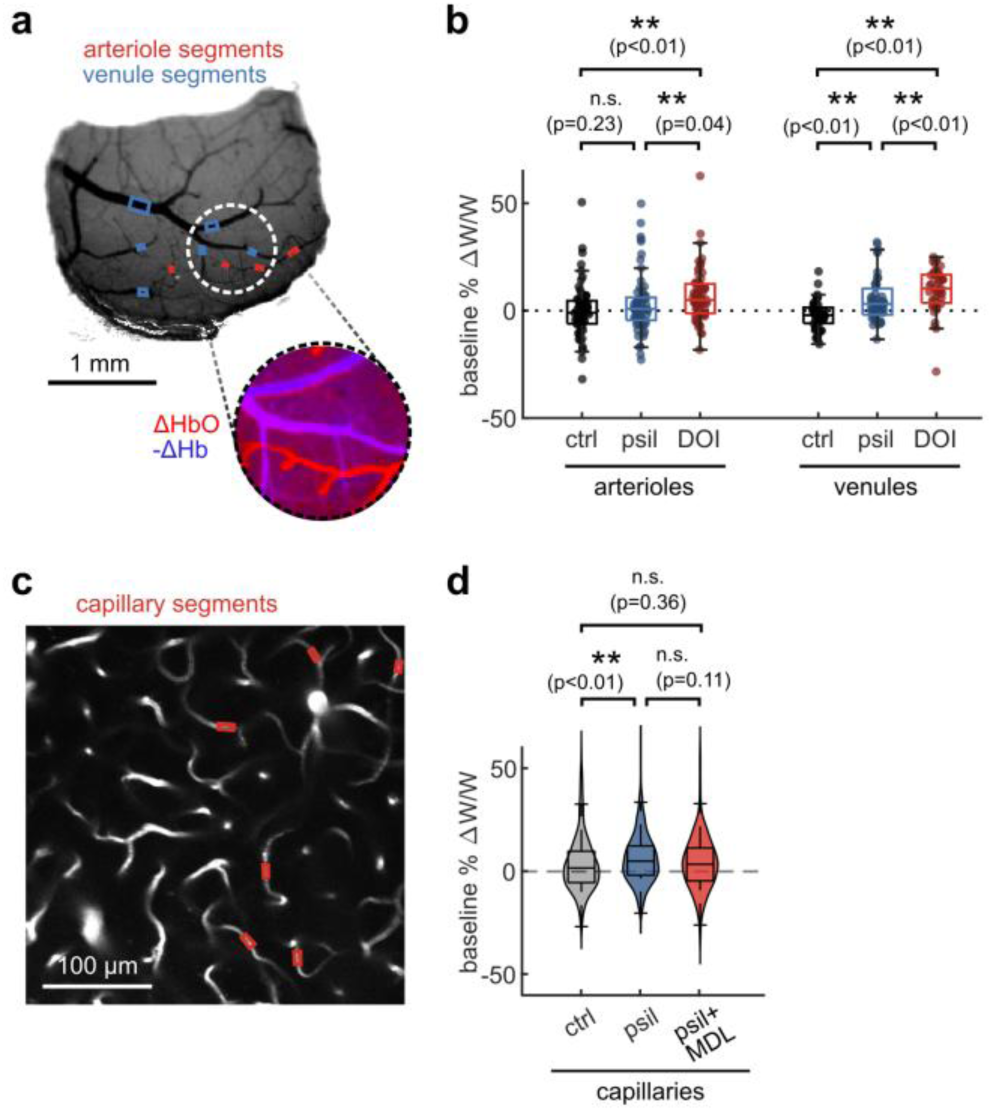
Increased baseline vessel width after psilocybin and DOI a: (top) Representative widefield 530 nm reflectance image for vessel classification and manual ROI drawing of arteriole and venule segments. (bottom) Oxygenation map used for vessel classification, calculated from the mean change in oxygenated hemoglobin during the stimulus period (in the red channel), which increases more in arterioles, and the inverse of the mean change in deoxygenated hemoglobin during the stimulus period (in the blue channel), which increases in venules. **b:** Relative change in baseline arteriole and venule diameter for control, psilocybin, and DOI treatment conditions. **c:** Representative two-photon image for manual ROI drawing for capillary segments. **d:** Relative change in baseline capillary diameter for control, psilocybin, and psilocybin+MDL treatments. All box plots display median, 25th, and 75th percentiles, with whiskers extending to the smallest/largest non-outliers. Significant effects between treatment conditions were determined with linear-mixed effects modeling followed by benjamini-hochberg correction for multiple comparisons (1 star: adj. p<0.05, 2 stars: adj. p<0.01).

### Prolonged NVC response in a whole-brain BOLD simulation led to misleading increases in functional connectivity measurements

To estimate the potential consequences that a prolonged NVC response could have on fMRI measurements, we performed numerical simulations of whole-brain BOLD signals. We used the Balloon-Windkessel NVC model, varying the parameters based on the empirically measured mouse blood flow responses before and after psilocybin. Since only two of the nine parameters of the model affect blood flow, only those parameters were tuned to fit the measured mouse responses (Fig. 6a). We first estimated the average stimulus-induced vessel flow velocity response by calculating a blood-volume-weighted average of the arteriole, venule, and capillary responses (Fig. 6b), then fit the model to the average flow response for the two conditions to simulate a pre and post-psilocybin BOLD response (Fig. 6c). Whole brain (80-region) simulations of neural activity (Fig. 6e) and the resulting BOLD signals in pre and post-psilocybin NVC conditions (Fig. 6f) revealed increased pairwise correlation of BOLD signals across nearly all brain area pairs (Fig. 6g-i), despite the fact that the underlying neural activity was identical. By simulating the NVC model over a larger space of blood flow parameters, we found a consistent gradient in mean BOLD-measured functional connectivity, with reduced or increased functional connectivity as the NVC response was shorter (pre-psilocybin) or longer (post-psilocybin), respectively (Fig. 6j)

**Fig. 6.**
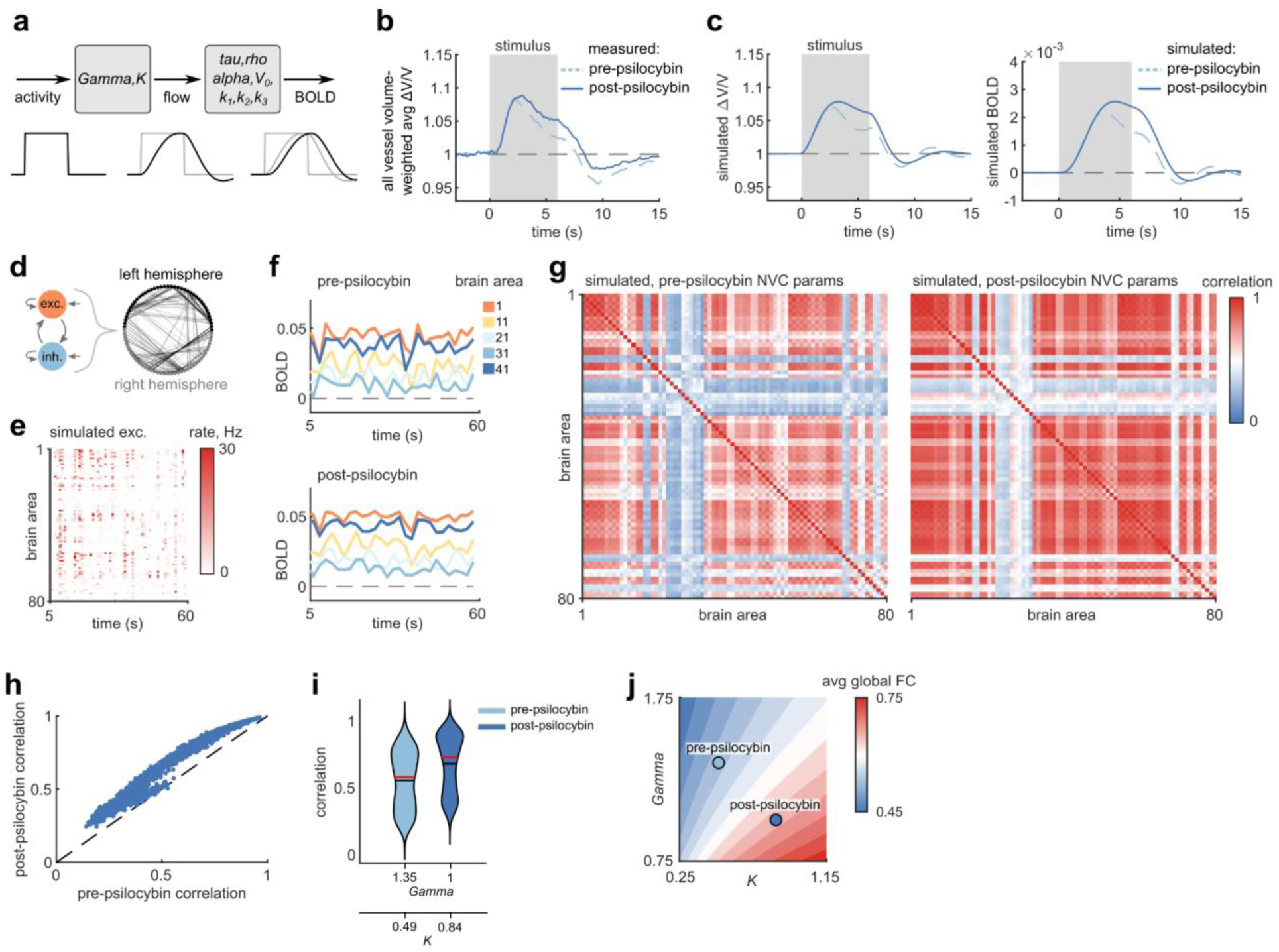
Prolonged NVC impacted BOLD measures of functional connectivity in whole-brain model a: Balloon-Windkessel model for simulating BOLD signals from neural activity, with parameters for simulating blood flow and BOLD indicated. **b:** Average change in stimulus-induced flow based on an estimated blood-volume weighted-average of measured arteriole, venule and capillary responses (56% capillaries, 22% arterioles, 22% venules)^41^ used in simulation. **c**: (left) Flow changes and (right) the simulated BOLD signals calculated with the Balloon-Windkessel model parameters best fitted to the pre- and post-psilocybin measurements in (b). **d:** (top) Overview of whole-brain simulation composed of 80 neural mass models (each with excitatory and inhibitory components) and their associated connectivity matrix. **e:** Raster plot of example simulated mean excitatory mean firing rates of all 80 brain areas. **f:** BOLD signals in a subset of brain areas activated as in (e), simulated based on fitted pre- (top) and post- (bottom) psilocybin model parameters. **g:** Average correlation matrices of 20 whole-brain BOLD simulations under pre (left) and post (right) psilocybin model parameters. **h:** Scatter plot of correlations between all brain area pairs, plotting pre vs post psilocybin conditions. **i:** Violin plots of distributions of brain area pairwise correlations for pre and post-psilocybin conditions (from data in h). The black and red lines display distribution mean and median, respectively. **j:** Average map of simulated global functional connectivity strength (mean of all pairwise correlations) from 20 simulations repeated across a 7×7 matrix of *Gamma* and *K* parameter values (the full simulated range is 0.25 - 1.75, in increments of 0.25), linearly interpolated between measured points. The fitted values of *Gamma* and *K* (from c) for pre- and post-psilocybin conditions values are shown.

## Discussion

Our findings show that psilocybin, at a psychoactive dose, alters neurovascular coupling in the awake mouse by prolonging the vascular response to visual stimulation without altering the underlying neuronal activity response. This decoupling specifically impaired the “recovery” phase of the response, as the initial peak increase after stimulation was largely unchanged after treatment. This could be due to a specific impact of psilocybin on the constriction phase of the vascular response – a critical component of the biphasic hemodynamic profile that typically follows stimulus-driven dilation – which might offer some clues to the mechanism of action. Under normal physiological conditions, vasodilation is primarily driven by excitatory pyramidal neurons through NMDA receptor–mediated release of prostaglandin E2 (PGE2) via COX-2 signaling, as well as by vasoactive interneurons, including VIP-positive and nNOS-positive neurons, which promote increased blood flow through vasoactive intestinal peptide (VIP) and nitric oxide (NO) signaling, respectively. In contrast, vasoconstriction can occur through multiple mechanisms, including activation of neuropeptide Y (NPY)-expressing interneurons ^39^, release of 20-HETE from astrocytes, and increased pericyte or vascular smooth muscle tone mediated by calcium-dependent signaling. These pathways help regulate or restore baseline blood flow following periods of neural activation.

Since psilocybin appeared to reduce this compensatory vasoconstriction, and that this effect was attenuated by pretreatment with MDL, 5-HT_2A_ receptor activation is implicated as a key mediator. However, psilocybin’s mechanism of action on NVC could involve much more than this single receptor. Other serotonin receptors with known roles in vasoactivity, including 5-HT_1A_, _1B_, and _2B_, are expressed across various cell types integral to vascular regulation, such as smooth muscle cells, pericytes, astrocytes, and endothelial cells ^32^; agonism at any of these sites could have direct or secondary effects on NVC.

While our study strongly implicates psilocybin in altering NVC dynamics, and given that this pattern was apparent across multiple measurement modalities, the limited scope of this work – focusing only on the mouse visual cortex – warrants caution in extrapolating these results too broadly. Given the known regional heterogeneity in 5-HT receptor expression ^40^, extending this analysis across multiple cortical and subcortical areas will be critical for understanding the generalizability of psilocybin’s effects on NVC. Additionally, our neural activity measurements were restricted to CaMKII-positive excitatory neurons, preventing us from characterizing potential psilocybin-induced changes in inhibitory interneuron and astrocyte responses, which may play critical roles in NVC regulation. Although we saw no change in activity in layer II/III neurons, it is possible that neurons in layer V – which contains many of the cortical neurons that express 5-HT_2A_ receptors – could show changes in visually evoked activity in response to psychedelics.

While acknowledging the limitations of our results, it is still useful to consider – if the pattern of psilocybin-induced NVC modulation observed in the mouse visual cortex holds true in human brain areas – what the implications may be in interpreting fMRI data in for psilocybin. Our whole-brain BOLD simulations show potential for misinterpretation of fMRI data toward increased functional connectivity across the brain. Regions may appear hyperactive or more functionally connected due solely to elevated or prolonged blood flow responses, even when neural activity remains unchanged. However, fMRI misinterpretation is not exclusive to a broadened NVC coupling response; if NVC is disrupted in a way that delays or dampens the vascular response, as in the case for DOI, this may lead to an underestimation of functional connectivity. The true nature of psilocybin’s effect on NVC may be more varied and complex: psilocybin may produce nonlinear dose-dependent effects, or differential effects by brain region. NVC modulation could be pronounced in regions with high levels of 5-HT_2A_ receptor expression such as the prefrontal cortex and visual cortex, while other subcortical areas may be less affected by psilocybin.

Regardless of the precise NVC modulation that psilocybin may have in humans, it is clear that consideration of vascular effects is crucial when interpreting neuroimaging data with psychedelics. To address these challenges, future human studies may consider combining fMRI with more direct neural recording methods to disentangle vascular and neural effects. Mapping the specific regional effects of psilocybin on NVC could enable analysis that can disentangle the NVC component to recover psychedelics’ effects on neural activity dynamics. The approach we have taken in this work – combining precise vascular measurements with neural recordings over multiple imaging modalities – provides a roadmap for understanding how NVC modulation may affect fMRI interpretation in the context of psychedelic research.

## Data Availability

Data documentation and smaller datasets are available at https://github.com/sn-lab/Psilocybin_NVC_Manuscript. Larger raw and processed imaging datasets are available upon request.

## Code Availability

Code for processing, analyzing, and plotting data and performing BOLD simulations are available online and described at https://github.com/sn-lab/Psilocybin_NVC_Manuscript.

## Acknowledgements

We thank Amanda Feilding for inspiring this study and for helpful discussions on the experimental design. This work was supported by NSF GRF (R.T.Z.), NIH grants R01MH128217 (A.C.K.) and R01MH137047 (A.C.K.), and One Mind – COMPASS Rising Star Award (A.C.K.).

## Author Contributions

R.T.Z., N.N., A.C.K. and C.B.S. conceived the project. R.T.Z. prepared mice, and conducted and analyzed 2PEF and widefield imaging data. M.I. analyzed 2PEF and widefield imaging data, performed statistical analyses, and conducted whole-brain BOLD simulations. R.T.Z. and C.L. performed surgeries. C.L., K.Y. and R.T.Z. performed habituation. M.Y. and K.Y. performed 2PEF volumetric imaging analysis. D.R. provided support for widefield imaging and data analysis. A.K. validated BOLD simulations. N.N., A.C.K., and C.B.S. provided supervision and edited the manuscript. R.T.Z. and M.I. wrote the initial version of the manuscript, and all authors contributed to the final version.

## Ethics declarations

A.C.K. has been a scientific advisor or consultant for Boehringer Ingelheim, Eli Lilly, Empyrean Neuroscience, Freedom Biosciences, and Xylo Bio. A.C.K. has received research support from Intra-Cellular Therapies. The other authors declare no competing interests.

## Methods

### Animals

All animal procedures complied with relevant ethical regulations and were performed after approval by the Institutional Animal Care and Use Committee of Cornell University (protocol number 2015-0029). All mice were housed in a climate-controlled facility kept at 22° C and 40-50% humidity, under a 12-hour light-dark cycle with ad libitum access to food and water. All mice were ∼5-7 weeks old WT (htr2a +/+) mice from Htr2a-flox transgenic lines. During 2PEF imaging experiments, mice were treated by intraperitoneal (IP) injection with either a control treatment (saline), psilocybin (1 mg/kg), or psilocybin after pretreatment with MDL100907 (1 mg/kg). During widefield imaging experiments, mice were treated by injection with either a control treatment (saline), psilocybin (1 mg/kg), or DOI (10 mg/kg).

### Surgery

Mice were anesthetized with isoflurane (3% for induction, 1% for maintenance) and placed on a feedback-controlled heating pad at 37° C. Surgeries were performed on a stereotaxic apparatus where the heads of mice were fixed with two ear bars. Ointment (Puralube, Dechra) was applied to both eyes for protection. Glycopyrrolate (0.002g/100g) was given intramuscularly to prevent fluid build-up in the lungs. Bupivacaine (0.125% ∼0.1mL) was administered below the scalp after being disinfected by iodine and 70% ethanol. An incision was made into the scalp to fully expose the skull. A 3-mm diameter craniotomy was made above V1 (A-P 3mm, M-L 2.5 mm from Bregma, centerline) on the right hemisphere. A 50 nL bolus of AAV1-CaMKII-GCaMP6f (Addgene) diluted to 10^12^ vg ml^-1^ was injected to the target V1 layer II/III (D-V -0.2 mm from the brain surface). A 3-mm diameter glass window then replaced the hole in the skull, and a titanium head plate was secured to the skull with Metabond (Parkell). Postoperative ketoprofen (5mg/kg) and dexamethasone (0.2 mg/kg) were administered subcutaneously, and the mouse was allowed to fully recover in a cage on a heating pad. Four weeks were allowed for viral expression before imaging.

### Habituation

To prepare for awake imaging, mice were habituated for 1-2 weeks prior to imaging. Imaging was performed with mice head fixed while awake with their bodies in a snug transparent tube. Habituation followed a progressively increased time spent head-fixed in the tube, starting at 5 minutes, and in increments of 10-15 minutes more each day working up to 1 hour of fixation. Mice were rewarded peanut butter during and after habituation.

### Visual stimulus

Visual stimulation experiments were performed in awake, head-fixed mice using a MouseGoggles Mono display positioned at the left eye, targeting the contralateral (right hemisphere) cortex for two-photon (2PEF) or widefield imaging. The display was aligned at 45° azimuth and 0° elevation relative to the mouse’s long axis. Visual stimuli consisted of a maximum-contrast blue square-wave grating with a spatial wavelength of ∼34.8°, extending across the entire display (covering 140° in azimuth and 122.5° in elevation of the visual field, due to the single clipped edge at the top of the circular display). The grating drifted at 1.5 Hz temporal frequency and was presented in four directions, each for 0.5 s in sequence, repeated three times for a total stimulus duration of 6 s per trial. Each trial lasted 16–24 s, with inter-trial intervals randomly selected between 10–18 s. The stimulus was repeated 25 times for 2PEF imaging and 15 for widefield imaging, due to data storage limitations of our widefield imaging setup.

### Two-photon excited fluorescence (2PEF) imaging

2PEF imaging was performed using a Ti:Sapphire laser (Coherent Vision S Chameleon; 80-MHz repetition rate, 75-fs pulse duration) at 920 nm to excite Texas Red-conjugated 70,000 MW Dextran and the CaMKII-GCaMP6f calcium indicator, with ∼35 mW power at the sample. A Zeiss 20x water immersion Objective (working distance = 2.3 mm; NA = 1) was used and heated by an objective heater set at 37° C. Imaging signals were acquired using ScanImage software (SI2022) into separate green- and red-colored channels (separated by a 560 nm long-pass dichroic; green channel using a 517/65 (center wavelength/bandwidth, both in nanometers) bandpass filter and red channel using 645/65. A TTL signal from the monocular display was acquired into an additional unused imaging channel for synchronizing imaging with visual stimuli.

For 2PEF imaging, mice were lightly anesthetized with isoflurane to perform a retro-orbital injection of Texas Red Dextran and were head-fixed to the awake imaging tube rig. After 30 minutes, mice were imaged for 30 minutes. This consisted of spontaneous and stimulated (25 trials, 18-24 s each) arbitrary line scan and frame scan imaging. During post-injection recording, 30 minutes after injection, mice were imaged for 30 minutes.

### 2PEF arbitrary line scan imaging

Arbitrary line scanning (Cycle rate = 166.67 Hz) was used to acquire blood flow velocity and diameter from vessel ROIs (n = 3 vessels/path), as well as calcium indicator fluorescence from neuron ROIs (n = 6 neurons/path). This was performed in layer II/III of the visual cortex at ∼180-200 µm depths. Preliminary imaging was performed to determine a location with high calcium fluorescence activation in response to the stimulus. ROIs were drawn using ScanImage ROI Group Editor, sequentially drawing ROIs to record vessel flow, neuron 1, neuron 2, vessel diameter, for three sets of 1 vessel and 2 neurons (Fig. 1b-c). A total of 12 ROIs were drawn with a pause between each to allow the scan to transition between lines. The duration of scanning for each ROI and pause was 0.25 ms, resulting in a single line scan path requiring 6 ms.

### 2PEF Arbitrary Line Scan Analysis

For analysis of two-photon arbitrary line scans in Matlab, logged data was first segmented into ROIs using the logged [x,y] positions (in degrees) of the scan mirror, a feature built into the ScanImage microscope control system. Since ScanImage logs the scan mirror positions using an auxiliary DAQ device that 1) samples data at a much slower rate than the PMT digitizer, and 2) skips four position samples at the end of each line scan to allow enough time to save the sampled data, the scan mirror positions were first resampled to the PMT data timestamps and had missing values filled by interpolation with an Akima spline (using the fillmissing() function in Matlab with the ‘makima’ method). This interpolation was manually checked for every imaging session to ensure accurate upsampling. Using the logged scan positions and the saved line ROI coordinates, the PMT data was segmented so that PMT values corresponding to scan positions near the drawn line ROIs were extracted for each ROI. Position values in degrees were converted into µm (0,0 located at the center of the image) using a conversion factor determined by imaging a calibration slide. Line scans taken with a high sample rate tend to have the scan mirror positions move faster in the middle of each line ROI and round off corners between ROIs, so samples taken across a line ROI are not linearly spaced. Since our method of calculating blood flow velocity from vessel line scans depend on linearly spaced sampling (detailed below), line scan segments were first resampled onto a linearly spaced position vector for every line ROI individually. Samples were excluded if they were acquired more than 4 µm away from the ROI (a distance selected so that all vessel line scan data were sampled inside the vessel walls), which often occurred near the ends of each line ROI where the scan mirror rounded off the corners to transition between ROIs.

To calculate blood flow velocity from linearized line scans down a vessel, line scans were formatted into 2D image frames where each row contains a linearized line scan and multiple rows represent successive line scans. Blood flow velocity can be observed in the 2D images by identifying the dark shadows caused by red blood cells travelling down the vessel, which appears as angled dark bands in the 2D image (with the slope of the dark streak proportional to the inverse of the flow velocity). The angle of dark bands (and hence blood flow velocity) was determined using a Radon transform based algorithm, as previously described ^42,43^. In brief, we perform background subtraction of the average projected data, window the line scan data into short time segments (50 lines) to accurately balance fast dynamics such as heart rate with slower responses, Radon transform the data, and then find the angle with maximal variance along distance, which is perpendicular to the streaks. This results in a vector of flow velocity, where each velocity measurement relates to the average velocity in that block of 50 line scans. Outliers in velocity measurements were manually defined using a minimum and maximum threshold.

To calculate calcium indicator fluorescence from linearized line scans across a neuron, the mean brightness of that line scan segment was calculated for every frame. To calculate the vessel width from linearized line scans across a vessel, a Gaussian function was fitted to the vessel cross section during every scan, and the vessel width was estimated as the full-width at half-maximum (FWHM) of the fitted Gaussian function. For vessels which were labelled brighter on the edges than in the center, a two-term Gaussian function was fitted where the two Gaussians traced the profile of each the two vessel edges, and the vessel width was estimated as the distance between the outer boundaries of the full-width at half-maximum (FWHM) of the two Gaussians.

Classification of neurons and vessels by responsivity (positively-responding, negatively-responding, or non-responding) was determined by first calculating the mean response at baseline (during the 3 s before visual stimulus onset) and during the 6-s stimulus (excluding the first 0.5 s transient period) for every stimulus repetition (n=25 repetitions) during the pre-injection time point: if a neuron or vessel had significantly greater mean response during the stimulus (Wilcoxon Sign Rank test, p<0.05), it was classified as a positively-responding; if it had significantly lower values during the stimulus, it was classified as negatively-responding; and if it had no significantly different values during the stimulus, it was classified as non-responding. To create mean calcium fluorescence change traces of positively-responding neurons (which were the majority of recorded neurons), all brightness vectors of positively-responding neurons which were recorded in both pre and post timepoints were first normalized to their respective baselines (F/F) and then had their baselines subtracted (ΔF/F). The mean and standard error of measurement (S.E.M.) were then calculated across all positively-responding neurons for each experimental condition individually. The mean response of each of these neurons were then calculated by averaging the response during the 0.5-6-s time period. Similarly, to create mean flow velocity and vessel width traces, vectors for all vessels that were recorded in both pre and post timepoints were normalized to their respective baselines (V/V for flow and W/W for width) and then had their baselines subtracted (ΔV/V and ΔW/W). The mean and standard error of measurement (S.E.M.) were then calculated across all vessels for each experimental condition individually. Since vascular responses tended to peak around 2.5 s after stimulus onset, the mean flow “rise” of these vessels were calculated by averaging the response during the 0.5-2.5-s time period, and the mean flow “recovery” was calculated by averaging the response during the 2.5-12-s time period.

### 2PEF Frame Scan Imaging

Frame scanning of stacks was done at 512×512 frames acquired at 1.06 Hz, with a step size of 1 µm. Laser power was manually increased during stack acquisition as fluorescence intensity weakened at greater tissue depths. The range of stacks included frames recorded from the brain surface – identified by surface vasculature and window interface – to 500 µm deep.

### 2PEF Frame Scan Analysis

Vessel stacks were analyzed in FIJI by acquiring 10-slice projections throughout the stack. 5-6 projections at evenly spaced intervals between 50-350 µm were selected. From these slices ROIs were manually drawn around 5-6 capillary segments (Fig 5c), to which a Gaussian function was fitted to obtain a FWHM which was used as the vessel width estimate.

### Widefield Multimodal Imaging

Widefield, multimodal imaging was performed by acquiring images using a Basler acA0240um-NIR enhanced camera. Interleaved images of 785-, 470-, 530-, and 565-nm illumination were acquired at 40Hz using a 785 nm laser (LD785-SEV300, ThorLabs), along with three LEDs with center wavelengths of 470 nm, 530 nm, 565 nm (ThorLabs) and filtered with the corresponding optical bandpass filters, respectively: 469/35nm, 532/10nm, and 560/10nm (ThorLabs). A 505 nm dichroic and 500 nm longpass filter separated the GCaMP fluorescence driven by 470 nm excitation for GCaMP imaging. A 4x 0.20NA objective (CFI Plan Apochromat Lambda D 4X, Nikon) was used. Imaging signals were acquired using a custom Matlab GUI. A TTL signal from the monocular display was recorded for synchronizing imaging with visual stimuli.

For widefield imaging, mice were lightly anesthetized with isoflurane and were head fixed in an awake imaging tube rig. After 30 minutes, mice were imaged for 30 minutes. This consisted of spontaneous and stimulated (15 trials, 18-24 s each) widefield imaging. During post-injection recording, 30 minutes after injection, mice were imaged for 30 minutes.

### Widefield multimodal analysis

Analysis of widefield multi-channel imaging videos was performed with custom Matlab scripts. All raw LED imaging channels and processed speckle images (see below) were Gaussian filtered using a 13×13 pixel window with a 2D Gaussian standard deviation of 3 pixels to reduce image noise. Raw and processed multi-channel imaging videos were used to calculate changes in blood flow speed, oxygenation, calcium indicator fluorescence, and vessel width (described below).

After these values were calculated (each on a frame-by-frame basis), time-series vectors of each measurement type were averaged over all 15 stimulus repetitions for each vessel segment and ROI type (parenchyma or GCaMP region) individually to get the average response vector for that vessel segment/ROI in that imaging session. The mean and standard error of measurement (S.E.M.) were then calculated across all vessel segments/ROIs for each experimental condition individually. The mean “rise” of these measurements were calculated by averaging the response during the 0.5-2.5-s time period, and the mean “recovery” was calculated by averaging the response during the 2.5-12-s time period. For calcium indicator fluorescence, the mean “response” was calculated by averaging the response during the 0.5-6-s time period.

#### Calculation of speed change

Laser speckle images (K) were converted from raw 785-nm excitation images by dividing the standard deviation (*σ)* by the mean intensity ⟨*I*⟩ of pixels within a 7×7 moving window:

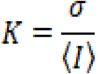

Inverse correlation time (*ICT*) was approximated from the speckle contrast *K* and speckle exposure time (*T*, 5 ms) using the simplified equation:^44^

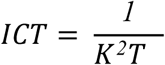

Changes in ICT values were calculated by normalizing all pixels in a frame to their pre-stimulus baseline average per trial (mean value during the 3-s pre-stimulus time period), making it equivalent to the normalized speed change.

#### Calculation of oxygenation change

Relative oxygenated and deoxygenated hemoglobin concentrations (Hb and HbO) were calculated using the 530-nm and 560-nm reflectance channels: First, the optical densities of every pixel in the 530-nm and 560-nm channels were calculated:

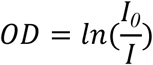

Where *I_0_* is the pixel value at baseline (calculated as the average value during the 3-s pre-stimulus period), *I* is the current pixel value, and *OD* is the optical density. Next, Hb and HbO concentrations were determined using the differential path lengths of the 530-nm and 560-nm channels and the expected extinction coefficients for Hb and HbO at these two wavelengths using the following formula and values:

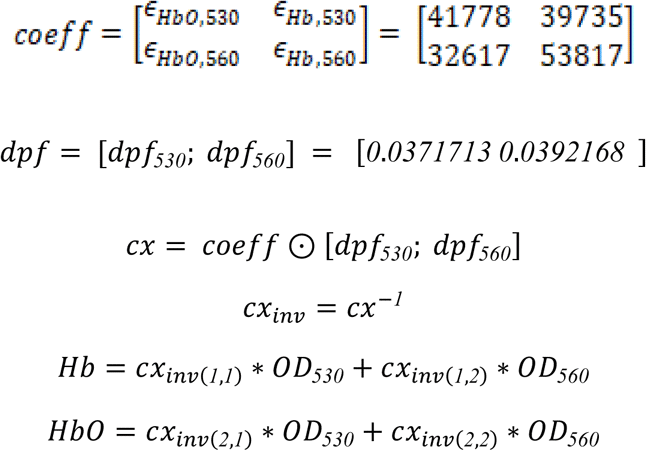

Where *coeff* is the extinction coefficient matrix, *dpf* is the differential pathlength factors, *cx* is the corrected coefficient matrix, and *cx_inv_* is the inverse of *cx.* HbT (total hemoglobin) was calculated as the sum of HbO and HbT concentrations.

#### Calculation of calcium indicator fluorescence change

To calculate calcium indicator fluorescence in the GCaMP-labelled tissue region, fluorescence was first corrected for changes in oxygenation. Fluctuations in hemoglobin concentration, particularly during functional hyperemia, introduce an optical artifact due to the strong absorption properties of oxy- and deoxyhemoglobin. As hemoglobin absorbs both excitation and emitted fluorescence light, local changes in blood volume and oxygenation can alter the detected fluorescence intensity independent of actual neural activity. Because of this, we performed oxy/deoxy absorption correction^55^ to account for these distortions impacting our GCaMP signal. The Hb and HbO values computed earlier were used to correct GCaMP fluorescence values using the following equations:

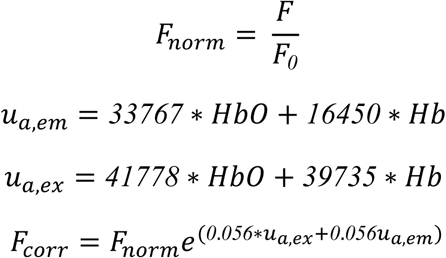

Where F is the raw fluorescence, F_0_ is the baseline fluorescence, F_norm_ is the normalized fluorescence, u_a,em_ is the absorption coefficient at the emission (530 nm) wavelength, u_a,ex_ is the absorption coefficient at the excitation (470 nm) wavelength, and F_corr_ is the corrected fluorescence.

#### Generating arteriole, venule, and parenchyma ROIs

To create ROIs for arterioles and venules, we first determined which vessels in the FOV were arterioles or venules using color-coded maps generated from the average Hb and HbO videos; these maps took advantage of the observation that the stimulus increases Hb more in arterioles than in venules, and that the stimulus causes a greater decrease in HbO in venules than in arterioles. The map of mean change in HbO from 0.5 – 6 s after stimulus onset was mapped onto the red channel, and the mean change in Hb was inverted and mapped onto the blue channel, creating a map of likely arterioles (in blue/purple) and venules (in red). This map, along with the increased branching and tortuosity of arterioles, was used to manually draw rectangular ROIs around vessel segments and classify these segments as arterioles or venules, where 3-7 segments of each type were labelled per imaging session. To determine the changes in the response measurements in arteriole and venule segments, binary image masks were created for every drawn rectangular vessel segment ROI, and the mean values inside the masks were calculated for every frame.

To create an ROI for the parenchyma, a binary image mask of the vessels was first created using image binarization using an adaptive threshold (imbinarize() function in Matlab) of the 530 nm reflectance channel, where vessels appear significantly darker than the surrounding tissue. The binary vessel mask was grown by 7 pixels (imdilate() in Matlab) to account for the 13×13 pixel window size of the Guassian filter, which blurs values up to 6 pixels in x/y directions, to minimize large value changes in the vessels from bleeding into the relatively smaller changes in parenchyma.

To create an ROI for the GCaMP virus-labelled parenchyma region, a binary image mask was first created around the brighter region of tissue in the 470 nm fluorescence channel. The mask was combined with the parenchyma mask to restrict the region to GCaMP-labelled parenchyma, excluding most vessels. All parenchyma and GCaMP-labelled region masks were manually inspected to ensure accuracy.

Using these vessel and parenchyma masks, all extracted values (Hb, HbO, HbT, ICT, GCaMP) were calculated on a frame-by-frame basis using their respective masks to create vectors of changes caused by the stimulus.

#### Calculation of vessel width

To calculate the width of the vessel segments within each manually-drawn rectangular vessel ROI, the center line of the vessel was calculated based on the angle of the drawn ROI and the image was rotated and collapsed along the vessel length to create a single average cross section vector. A Gaussian function was then fit to the cross section, and the vessel width was estimated as the full-width at half-maximum (FWHM) of the fitted Gaussian function. This process was performed on a frame-by-frame basis to create vectors of width changes caused by the stimulus.

### Statistical analysis

Significant differences between all mean value changes for different experimental conditions were determined using custom Matlab scripts for linear mixed effects (LME) modelling using the fitlme() function. The general form of the equation used was:

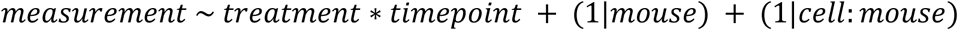

Where *measurement* is the continuous variable being evaluated (e.g. mean dF/F response; mean “rise” dV/V; mean “rise” dV/V), treatment is the treatment category (control or treatment), timepoint is the session category (pre or post), mouse is the categorical mouse ID (e.g. mouse 1-32), and cell is the categorical vessel/neuron ID (e.g. neuron 1-3; vessel 1-6). P-values for the treatment effect were obtained from the post:treatment entry from the ANOVA table. For each common set of statistical comparisons (e.g. 2PEF control vs psilocybin, Fig. 1 and Extended Data Fig. 1), the set of p-values collected were corrected for multiple comparisons using the Benjamini-Hochberg procedure, using the fdr_bh() Matlab function ^45^; all p-values reported in this study are adjusted p-values after multiple comparison correction.

### Balloon-Windkessel model and whole-brain simulations

The Balloon-Windkessel model was simulated in python and in custom Matlab scripts using the neurolib python simulation framework ^46^, modified to accept custom NVC parameters as function inputs and output both the final simulated BOLD signal as well as the blood flow internal state variable. The starting point of the model parameters were based on the set of parameters previously fitted to human fMRI data ^47^.

To fit model parameters to our measured mouse blood flow response in pre and post-psilocybin conditions, we first estimated the average whole-brain vessel flow response in these two conditions by taking a weighted average of the capillary responses (from Fig. 1i), arteriole responses, and venule responses (from Fig. 2g). These responses were weighted by estimated blood volume flowing through each vessel type in the mouse brain: 56% in capillaries, 22% in arteries/arterioles, and 22% in veins/venules ^41^. The model parameters were then fit based on least-squares error of the average measured all-vessel flow response to the simulated flow response of an adaptive linear-nonlinear (ALN) neural mass model to a 6-s square wave excitation input.

Whole-brain simulation was performed using the neurolib framework in python using the included “gw” 80-region DTI/fMRI dataset. Each of the 80 nodes was simulated as an ALN neural mass model with excitatory and inhibitory components, where each of the nodes were connected based on the connectivity and delay matrices from the gw dataset. To compare whole-brain BOLD signals resulting from pre and post-psilocybin Balloon-Windkessel model parameter conditions, A total of 20 1-minute whole-brain simulations were performed, each simulation repeated under the pre-psilocybin and post-psilocybin conditions, where each simulation pair contained identical simulated neural activity patterns. To obtain whole-brain BOLD signals across a wider range of model parameters, the Gamma and K parameters were individually varied from 0.25 to 1.75 (in increments of 0.25) and the same set of 20 simulations were repeated for each unique set of parameters.

To calculate functional connectivity (FC) from simulated whole-brain BOLD signals, the Pearson correlation was obtained for each pair of the 80 brain regions and mapped onto an FC matrix. To calculate the global FC strength from the FC matrix, the mean of the upper triangle (only including unique pairs of different brain regions) of the FC matrix was calculated.

---

**Extended Data Fig. 1, related to Fig. 1.**
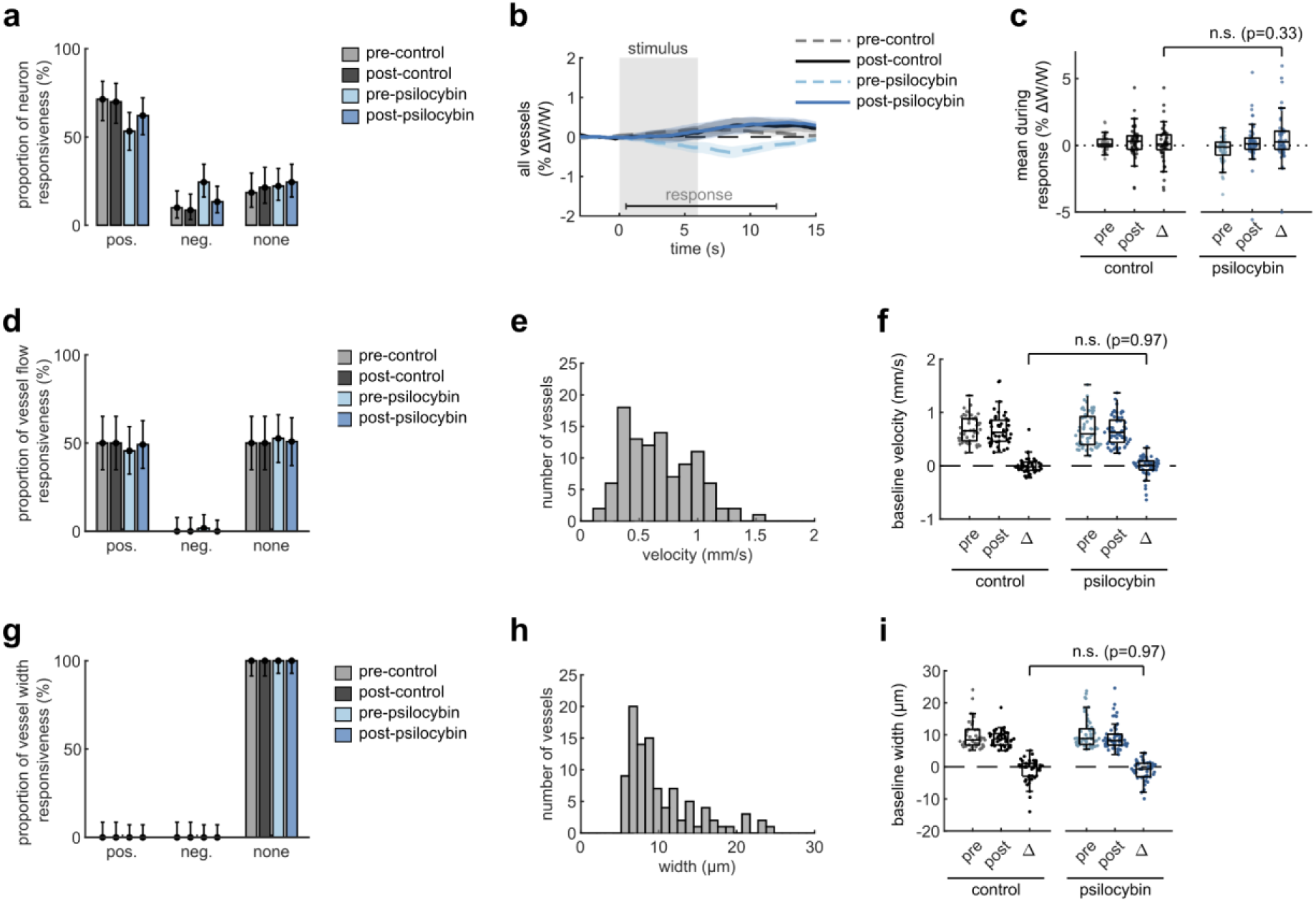
Effects of Psilocybin on V1 capillary dilation, neuron and vessel responsiveness, and baseline vessel flow. **a:** Bar plots of neuron classification by response type: positively responding neurons with significantly increased fluorescence during the stimulus period (0.5-6 s); negatively responding neurons with increased fluorescence after the stimulus offset (6.5-12 s); and neurons with no significant response. **b:** Mean vessel diameter change of all vessels for each treatment condition. **c:** Box plots of mean vessel diameter change during the response period (0.5-12 s) of each vessel in all conditions, with the mean post-pre difference values (Δ) shown for each treatment. **d:** Bar plots similar to (c), for vessel flow response type during the stimulus period (0.5-6 s). **e:** Histogram of baseline flow velocity across the capillaries measured. **f:** Box plots of baseline flow velocity in capillaries for all treatment conditions, as well as post-pre differences. **g-i**: Bar plots (g), histogram (h), and box plots (i) similar to (d-f) but for vessel diameter responsiveness and baseline vessel diameters. All mean traces are shown with +/-S.E.M. in shaded regions. Box plots display median, 25th, and 75th percentiles, with whiskers extending to the smallest/largest non-outliers. Significant effects between treatment conditions were determined with linear-mixed effects modeling followed by benjamini-hochberg correction for multiple comparisons (1 star: adj. p<0.05, 2 stars: adj. p<0.01).

**Extended Data Fig. 2, related to Fig. 2.**
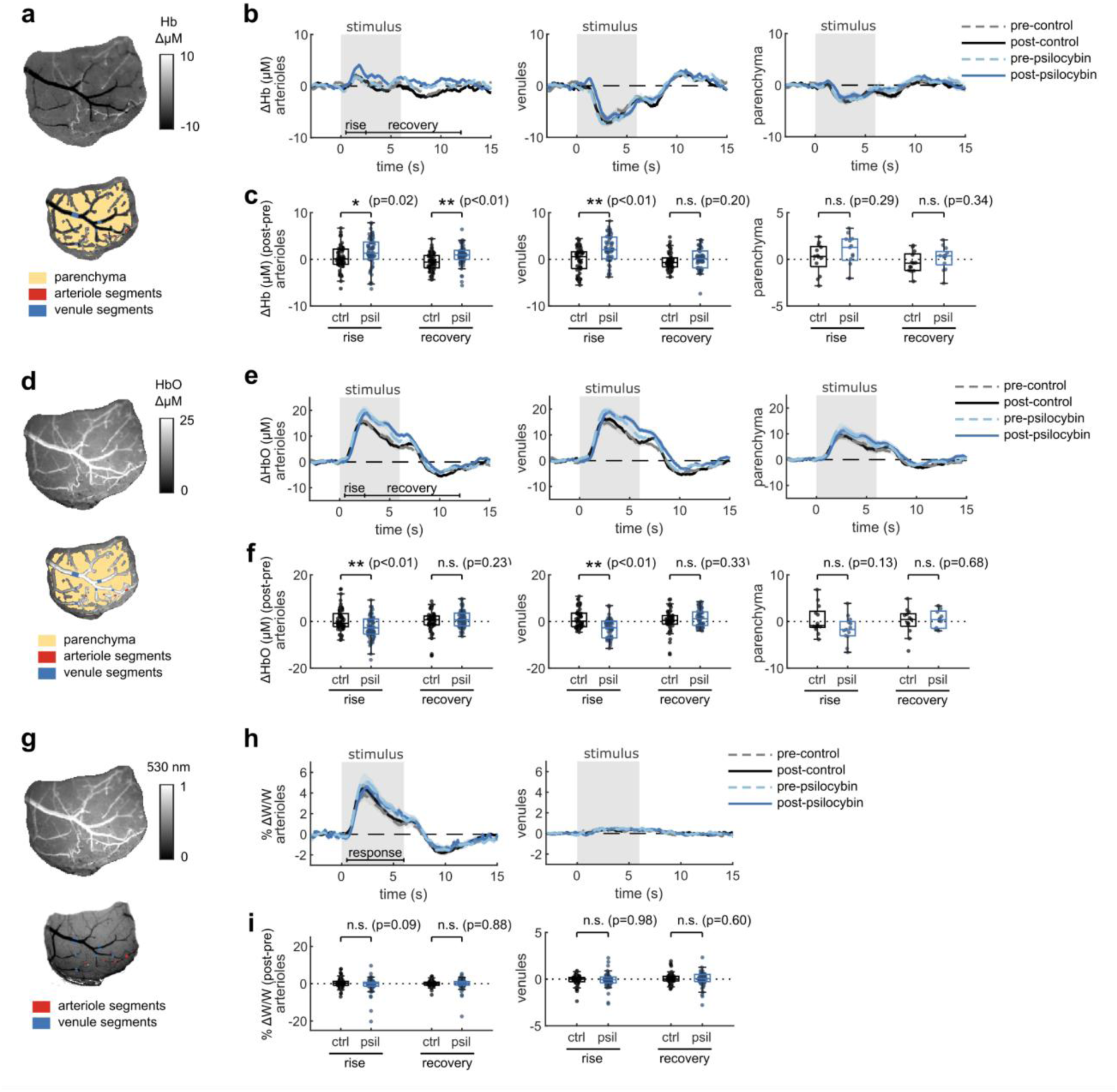
The effects of psilocybin on widefield measurements of oxygenation and vessel dilation. Same figure structure, labeling, and statistical analysis as in Figure 2 (f-h), but for deoxygenated hemoglobin (Hb) (a-c), oxygenated hemoglobin (HbO) (d-f), and vessel diameter (g-i).

**Extended Data Fig. 3, related to Fig. 3.**
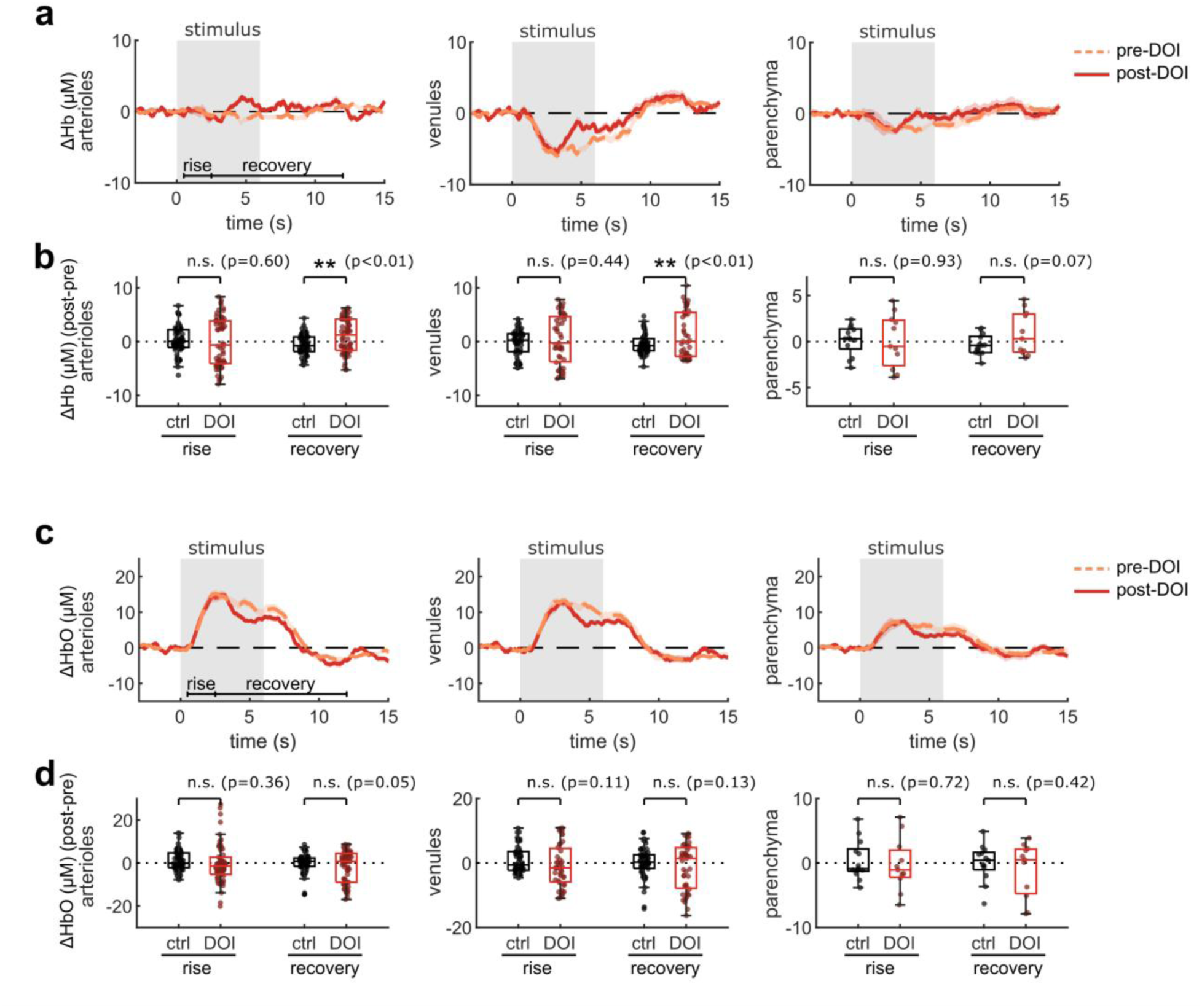
Effects of DOI on widefield measurements of oxygenation. Same figure structure, labeling, and statistical analysis as in Figure 3 (e-h), but for deoxygenated hemoglobin (Hb) (a-b) and oxygenated hemoglobin (HbO) (c-d).

**Extended Data Fig. 4, related to Fig. 4.**
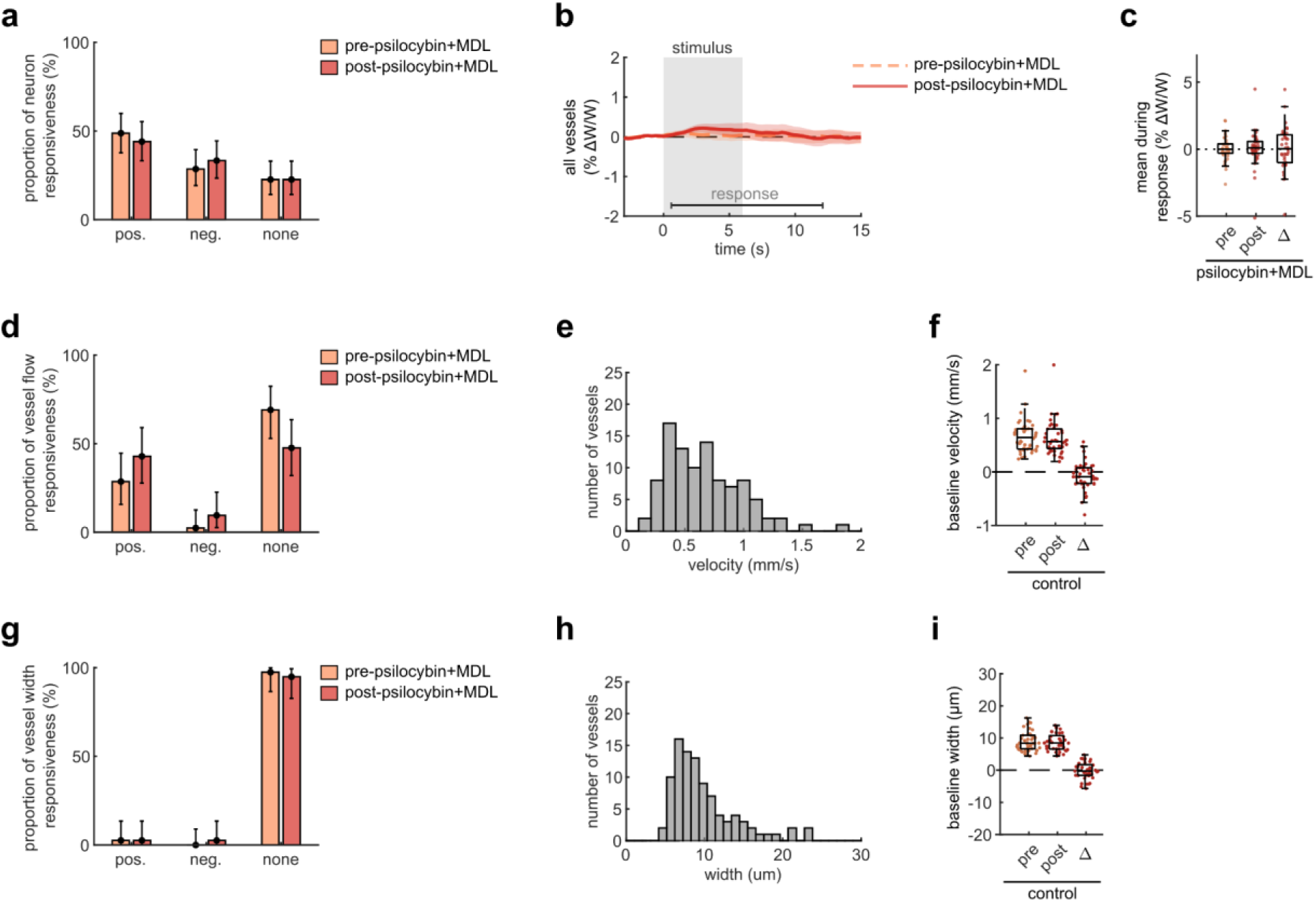
Effects of Psilocybin and MDL on V1 capillary dilation, neuron and vessel responsiveness, and baseline vessel flow. Same figure structure, labeling, and statistical analysis as in Extended Data Fig. 1, but for the psilocybin with MDL pretreatment condition.

